# Functional dissection of metabolic trait-associated gene regulation in steady state and stimulated human skeletal muscle cells

**DOI:** 10.1101/2024.11.28.625886

**Authors:** Kirsten Nishino, Gabriella Bloss, Jacob O. Kitzman, Stephen C.J. Parker, Adelaide Tovar

## Abstract

Type 2 diabetes (T2D) is a common metabolic disorder characterized by dysregulation of glucose metabolism. Genome-wide association studies have defined hundreds of signals associated with T2D and related metabolic traits, predominantly in noncoding regions. While pancreatic islets have been a focal point given their central role in insulin production and glucose homeostasis, other metabolic tissues, including liver, adipose, and skeletal muscle, also contribute to T2D pathogenesis and risk. Here, we examined context-specific genetic regulation under basal and stimulated states. Using LHCN-M2 human skeletal muscle cells, we generated transcriptomic profiles and characterized regulatory activity of 327 metabolic trait-associated variants via a massively parallel reporter assay (MPRA). To identify condition-specific effects, we compared four different conditions: (1) undifferentiated, or (2) differentiated with basal media, (3) media supplemented with the AMP analog AICAR (to simulate exercise) or (4) media containing sodium palmitate (to induce insulin resistance). RNA-seq revealed these treatments extensively perturbed transcriptional regulation, with 498-3,686 genes showing significant differential expression between pairs of conditions. Among differentially expressed genes, we observed enrichment of relevant biological pathways including muscle differentiation (undifferentiated vs. differentiated), oxidoreductase activity (differentiated vs. AICAR), and glycogen binding (differentiated vs. palmitate). The results of our MPRA found broadly different levels of activity between all conditions. Our MPRA screen revealed a shared set of 7 variants with significant allelic activity across all conditions, along with a proportional number of variants showing condition-specific allelic bias and the total number of active oligos per condition. We found that a lead variant for serum triglyceride levels, rs490972, overlaps SP transcription factor motifs and has differential regulatory activity between conditions. Comparison of MPRA activity with paired gene expression data allowed us to predict that regulatory activity at this locus is mediated by SP1 transcription factor binding. While several of the MPRA variants have been previously characterized in other metabolic tissues, none have been studied in these stimulated states. Together, this work uncovers context-dependent transcriptomic and regulatory dynamics of T2D- and metabolic trait-associated variants in skeletal muscle cells, offering new insights into their functional roles in metabolic processes.

## Introduction

Type 2 diabetes (T2D) is a common metabolic disorder characterized by dysregulation of glucose metabolism and is a major public health issue, affecting over 483 million individuals worldwide (90% of 537 million diabetes mellitus cases as of 2021) (IDF, 2021). T2D is a complex disease and results from a combination of both genetic and environmental factors. Previous genome-wide association studies (GWAS) have successfully mapped over 1,000 independent signals to roughly 600 loci in the genome (Suzuki et al. 2024; Mahajan et al. 2022; Vujkovic et al. 2020). Because the majority of these signals reside far from genes in noncoding regions, progress to translate them to specific disease mechanisms has proceeded slowly. This is due to several challenges including difficulty predicting the functional consequences of noncoding variants, tight linkage between neighboring genetic variants, and lack of certainty about which cell types and environmental contexts are casual to the disease (Cano-Gamez and Trynka 2020).

T2D research has focused on pancreatic islets due to their direct connection to glucose regulation by insulin secretion; however, metabolic disease pathogenesis and risk is distributed across other important peripheral metabolic tissues, including the liver, adipose, and, the tissue of focus for this study, skeletal muscle. Glucose levels are regulated in skeletal muscle through both insulin-independent and insulin-dependent mechanisms, modulating delivery, transport, and metabolism (Hulett et al. 2022). Skeletal muscle is responsible for 80% of postprandial glucose uptake, and therefore is a tissue of high interest when studying metabolic disorders (DeFronzo and Tripathy 2009). Previous studies have used functional genomic data collected from primary skeletal muscle samples to link T2D GWAS signals and target genes (Scott et al. 2016; Orchard et al. 2021; Varshney et al. 2023), but few studies have further probed context-specific effects of these variants. Exploring the behavior of variants across stimulatory states is a crucial next step to reveal how dynamic environmental changes and the associated shifts in cell states affect the functional impact of disease-associated variants.

Here, we used the human skeletal muscle cell line LHCN-M2 (Zhu et al. 2007) across basal and stimulatory states to investigate cellular environmental effects on regulatory features of T2D and metabolic trait-associated variants (**Figure 1**). To examine developmental state changes, we differentiated LHCN-M2 proliferating myoblasts into terminally differentiated myotubes. To emulate different environmental stressors, we subsequently perturbed myotubes with either the AMP analog 5-aminoimidazole-4-carboxamide ribonucleotide (AICAR) to induce oxidative stress (Dolinar et al. 2018) and mimic exercise, or palmitate, a fatty acid known to cause insulin resistance (Mäkinen et al. 2017). We examined these stimulations’ effects on gene expression by performing differential expression analysis using bulk RNA-seq data from basal and perturbed cells. Then, to assess regulatory activity across a library of 327 metabolic trait-associated variants, we constructed and implemented a massively parallel reporter assay (MPRA) under the same conditions. Our study builds upon the current understanding of how metabolic disease-associated regulatory elements may be influenced by the cellular environment.

**Figure 1.**
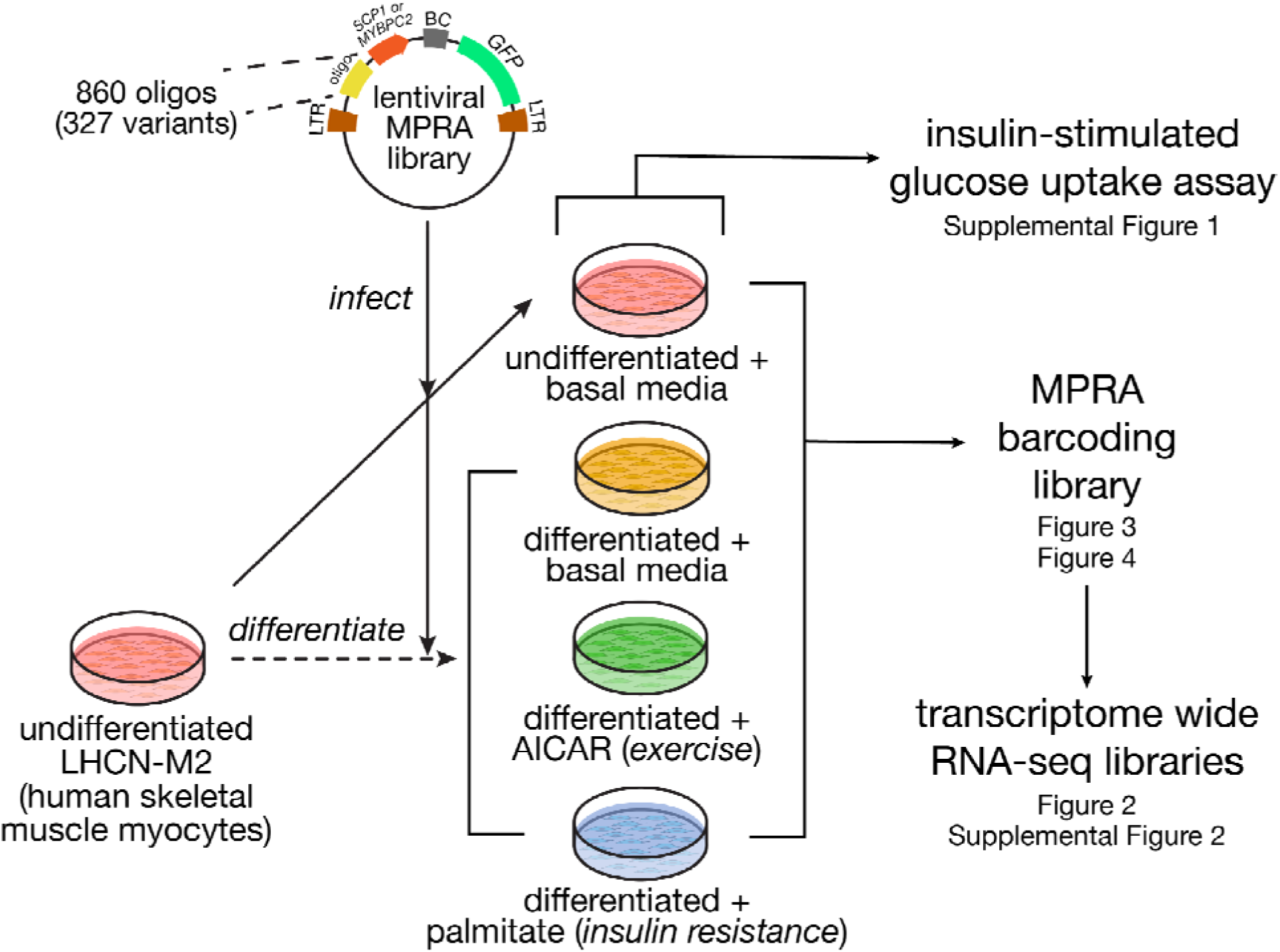
Study overview. We synthesized a library of 860 oligos spanning 327 variants with either the super core (SCP1) or muscle-specific (MYBPC2) promoter. This library was used to infect undifferentiated LHCN-M2 human skeletal muscle myocytes, and subsequently, a portion of the cells were differentiated into myotubes. These were stimulated with AICAR to mimic exercise or palmitate to induce insulin resistance, or left untreated. Across these four conditions, we measured insulin-stimulated glucose uptake, gene expression (by bulk RNA-seq) and variant regulatory activity (by MPRA).

## Results

### Differentially expressed genes highlight developmental biology and cellular context

To model stimulation states relevant to metabolic disease, we cultured LHCN-M2 skeletal muscle cells in four conditions: undifferentiated with basal media, differentiated with basal media, differentiated with AICAR-supplemented media, and differentiated with palmitate-supplemented media. We then examined insulin-stimulated glucose uptake, a physiological endpoint reflective of relevant muscle biology. We compared the fold-change in glucose uptake with or without insulin stimulation across conditions and observed a significant reduction in insulin-stimulated uptake in differentiated cells treated with palmitate relative to cells in basal media in both differentiation states (**Supplemental Figure 1**; *p* = 0.016 for undifferentiated, *p* = 0.016 for differentiated basal; Wilcoxon signed-rank test). Palmitate is commonly used to induce insulin resistance in experimental models, and these results are consistent with prior observations that palmitate exposure diminishes responsiveness to insulin in skeletal muscle cells (Mäkinen et al. 2017; Coll et al. 2008; Tokarz et al. 2023). By contrast, insulin-stimulated uptake did not differ from the undifferentiated condition for differentiated cells with basal and AICAR-supplemented media (*p =* 0.556 and *p =* 0.067, respectively). Finally, differentiated cells with AICAR-supplemented media showed a modest reduction in uptake compared to the differentiated cells with basal media (*p* = 0.067). This is likely due to increased baseline glucose uptake in the former group compared to the latter, which reduces the relative effect of insulin addition (**Supplemental Table 1**).

We performed bulk RNA-seq under the same conditions in quadruplicate, and clustered the resulting samples by principal component analysis (**Figure 2A, Supplemental Table 2**). Differentiation state was strongly separated along PC1 (54.40% of variance). The conditions using differentiated myotubes were further separated along PC2 (30.89% of variance) with AICAR-stimulated samples showing the greatest separation. We first compared the undifferentiated and basal differentiated conditions, and found 3,130 differentially expressed genes with DESeq2 (**Figure 2B**, **Supplemental Figure 2A**, **Supplemental Table 3**; log_2_FC > 1, FDR < 5%) (Love et al. 2014). Among the strongest changes was *MYF5*, a key myogenic fate determination factor which was highly downregulated (log_2_FC ∼ -5; FDR-adjusted *p* < 10^-300^), consistent with its known expression pattern *in vivo* (Arnold and Braun 1999). Conversely, *MYH3* was strongly induced during differentiation (log_2_FC ∼ 5.22; FDR-adjusted *p* < 10^-300^). Enrichment analysis highlighted pathways involved in cell growth and division, such as DNA metabolism, cell division cycle, and ribosome biogenesis as highly expressed in undifferentiated samples (**Figure 2C**, **Supplemental Table 4**), consistent with myoblasts’ role as proliferating precursors of mature skeletal muscle cells (Vaughan and Lamia 2019), and with growth arrest due to permanent withdrawal from the cell cycle during differentiation (Myers et al. 2004). Conversely, pathways upregulated during differentiation included known in muscle cell development and morphogenesis gene sets, and actin filament based processes involved in myoblast elongation (Lehka and Rędowicz 2020). Taken together, these results highlight extensive transcriptional dynamics that direct the transition from proliferative myoblasts to specialized, mature myotubes.

**Figure 2.**
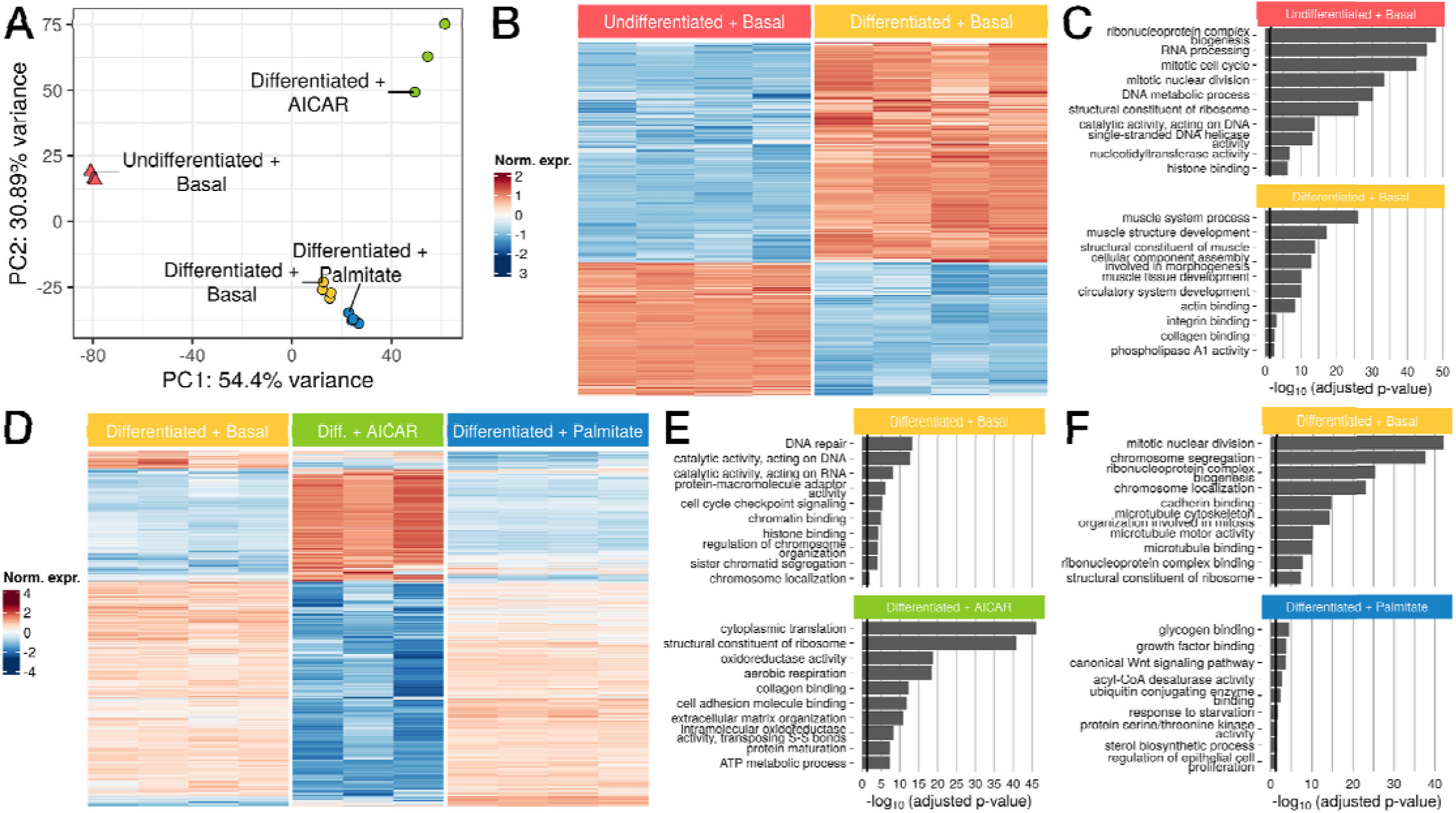
Gene expression varies widely after myoblast to myotube differentiation and in response to AICAR and palmitate. **(A)** Principal components analysis (PCA) of samples by gene expressi n. **(B)** Heatmap of genes differentially expressed between undifferentiated and differentiated samples (n=3,130) **(C)** Upregulated GO terms enriched in undifferentiated (top) or differentiated samples (bottom). **(D)** Heatmap of genes differentially expressed between differentiated LHCN-M2 myotubes in basal vs AICAR or palmitate-stimulated conditions (n=3,570) **(E,F)** Upregulated GO terms enriched in (**E**) differentiated basal (top) or AICAR-stimulated samples (bottom) and in **(F)** basal (top) or palmitate-stimulated samples (bottom). Black lines in **(C)**, **(E)**, and **(F)** correspond to *p* = 0.05.

To evaluate stimulus-specific effects, we performed pairwise comparisons between each perturbation (AICAR, palmitate) and the basal differentiated condition. First, we identified 3,686 differentially expressed genes between the AICAR-stimulated samples and differentiated samples (**Figure 2D**, **Supplemental Figure 2B**, **Supplemental Table 5**; log_2_FC > 1, FDR < 5%). In AICAR-stimulated cells, we observed a significant increase in *MYC* expression (log_2_FC ∼ 2.58; DESeq2 FDR-adjusted *p* = 1.757 x 10^-207^). Several studies have reported significant increases in *MYC* expression after both resistance and endurance exercise in humans and mouse models (Jones et al. 2023; Broholm et al. 2011; Trenerry et al. 2007; Popov et al. 2019). By contrast, *SIX1* was highly downregulated in AICAR-stimulated cells (log_2_FC ∼ -2.13; DESeq2 FDR-adjusted *p* = 1.138 x 10^-104^). Previous studies have reported decreased *SIX1* expression and activity in response to skeletal muscle overload (Gordon et al. 2012; Ramachandran et al. 2019; Kostek et al. 2007). Moreover, *SIX1* is an important developmental transcription factor that regulates skeletal muscle differentiation and promotes myoblast fusion to generate myotubes (Niro et al. 2010; Bessarab et al. 2008; Ridgeway and Skerjanc 2001; Fougerousse et al. 2002). Pathway enrichment analysis identified DNA repair and cell cycle-associated genes as significantly downregulated following AICAR treatment (**Figure 2E**, **Supplemental Table 6**), consistent with previous studies showing that AICAR inhibits proliferation and promotes cell cycle arrest (Nagata et al. 2004; Igata et al. 2005). In contrast, genes more highly expressed in AICAR-perturbed cells were related to protein synthesis and cellular oxidative metabolism, with pathways such as cytoplasmic translation, maintenance of ribosome structure, and oxidoreductase activity. AICAR-perturbed cells likely undergo adaptive processes such as translation of stress-response genes and preservation of ribosome structures to combat the effects of oxidative damage (Shcherbik and Pestov 2019). Oxidoreductase activity is responsible for maintaining redox homeostasis, and introduction of oxidative stress by AICAR would cause an imbalance of reactive oxygen species, therefore requiring higher levels of redox activity (Sies et al. 2017).

Response to palmitate identified 498 differentially expressed genes versus basal media (**Figure 2D**, **Supplemental Figure 2C**, **Supplemental Table 7**; log_2_FC > 1, FDR < 5%). Several lipid-associated genes were highly upregulated in palmitate-stimulated samples including *SCD* (log_2_FC ∼ 1.66; DESeq2 FDR-adjusted *p* = 6.329 x 10^-95^) (Voss et al. 2005; Dziewulska et al. 2020)*, PLIN2* (log_2_FC ∼ 1.97; DESeq2 FDR-adjusted *p* = 3.475 x 10^-49^) (Daemen et al. 2018; Cho and Kang 2015), *FADS1* (log_2_FC ∼ 1.18; DESeq2 FDR-adjusted *p* = 1.98 x 10^-48^) (Karjalainen et al. 2024; Selvaraj et al. 2022)*, and ANGPTL4* (log_2_FC ∼ 4.55; DESeq2 FDR-adjusted *p* = 4.663 x 10^-45^) (Kuo et al. 2024; McCulloch et al. 2020), all of which are implicated in insulin resistance. Significantly downregulated pathways after palmitate treatment were related to mitotic processes such as nuclear division and chromosome segregation (**Figure 2F**, **Supplemental Table 8**). While this appears to contradict the growth arrest expected in differentiated myotubes, palmitate is known to further inhibit proliferation and induce differentiation (Grabiec et al. 2015). Genes highly expressed in palmitate-stimulated cells were associated with glycogen binding, growth factor binding and the Wnt signaling pathway. Increases in circulating fatty acids such as palmitate promote insulin resistance and modulate glycogen binding and synthesis (Samuel and Shulman 2016; Roden et al. 1996; Dimopoulos et al. 2006). Additionally, recent work implicates the Wnt signaling pathway as a mediator of insulin resistance (Suren Garg et al. 2023). Altogether, these results highlight the unique context-specific pathways that are enriched for each condition, demonstrating the importance of developmental and environmental conditions on gene expression.

### Massively parallel reporter assay identifies allelic activity of oligos

We next used an MPRA to investigate the regulatory basis of the transcriptional differences between these stimulatory states. We constructed a library of 860 200-bp oligos, spanning 327 variants, including metabolic trait-associated variants that were previously characterized in luciferase assays (Orchard et al. 2021; Khetan et al. 2021; Roman et al. 2015; Pandey et al. 2024; Sinnott-Armstrong et al. 2021) and variants in tight linkage disequilibrium to measures of physical activity (Wang et al. 2022). To compare generic and tissue-specific promoter contexts, we cloned this oligo library into an MPRA vector with either the SCP1 synthetic housekeeping promoter or the promoter for the muscle-specific gene *MYBPC2*. Each cloned oligo fragment was tagged with a short randomized 20mer barcode such that each oligo could be associated with multiple barcodes (median = 927 barcodes/oligo) to provide internal replication. Subsequently, we delivered this library to LHCN-M2 human skeletal muscle cells by stable lentiviral transduction under all four conditions as mentioned above. After delivery, we deeply sequenced these MPRA barcodes from mRNA and genomic DNA (gDNA), with the ratio of these counts serving as a measure of each oligo’s activity in the MPRA.

After filtering to oligos with at least 10 unique barcodes, we recovered 735 oligos (85.5% of total) spanning 295 variants (90.2% of total). Further separating these data by promoter context, we recovered 720 oligos (83.7%) spanning 294 variants (89.9%) paired with the MYBPC2 promoter and 721 oligos (83.8%) spanning 290 variants (88.7%) paired with the SCP1 promoter. Across the two promoters, we recovered 700 oligos (81.4%) in common spanning 288 variants (88.1%). Notably, the results of clustering on MPRA signal (**Figure 3A**) mirrored the clustering based upon bulk RNA-seq (**Figure 2A**), with the differentiation and treatment separating samples on the first two principal components (42.4% and 12.8% of variance, respectively). As with RNA-seq PCA, these differences were not driven by read coverage or barcode recovery (**Supplemental Table 9**). A similar grouping was observed by PCA restricted to oligos paired with the *MYBPC2* promoter (**Supplemental Figure 3A**) or those paired with the SCP1 promoter (**Supplemental Figure 3B**), indicating that promoter context does not drive the clustering.

**Figure 3.**
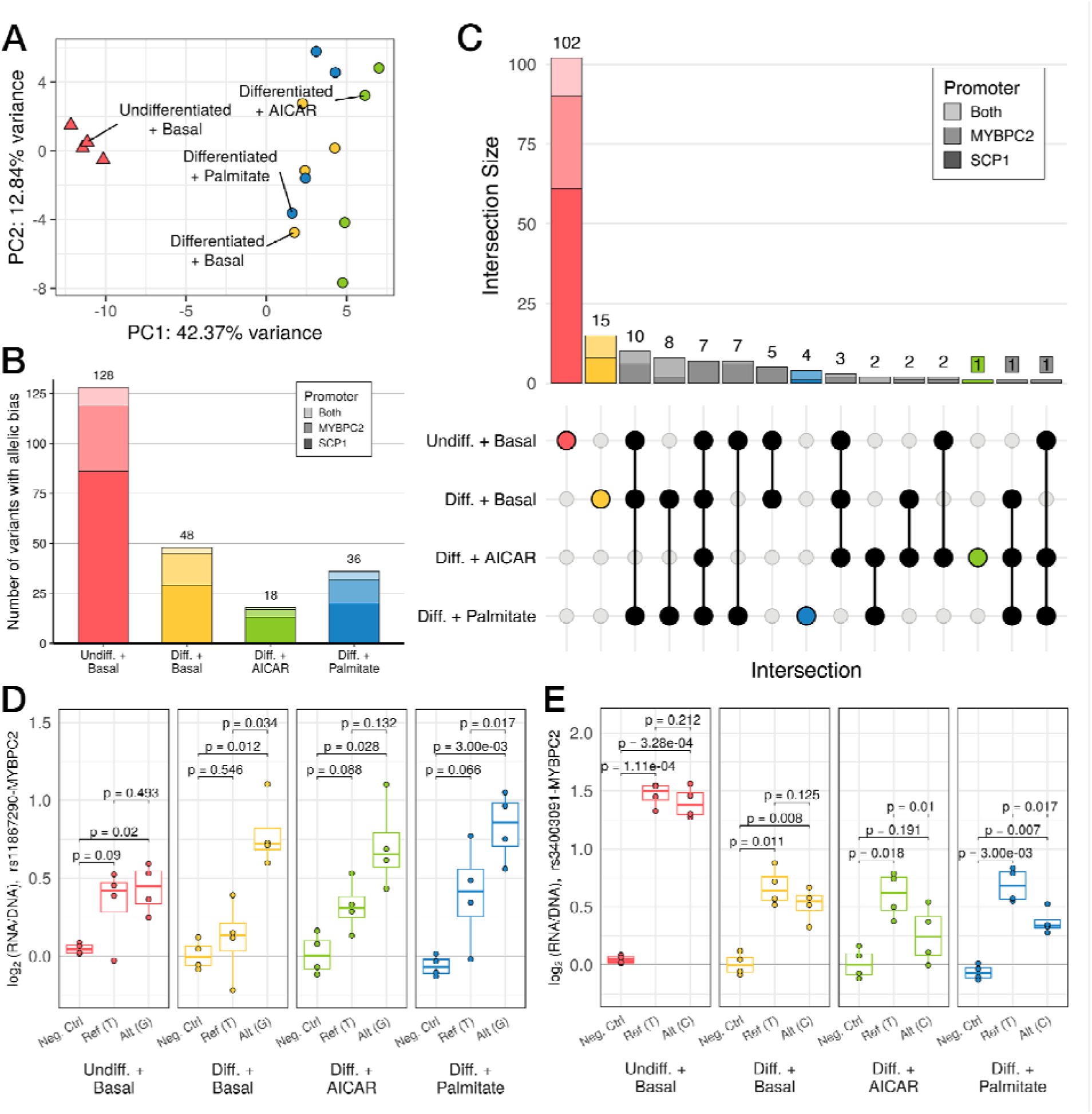
Variation in regulatory activity measured by MPRA recapitulates whole-transcriptome patterns. **(A)** Principal components analysis (PCA) of MPRA activity across 860 oligos. **(B)** Stacked bar plot displaying the number of variants that display differences in regulatory activity across both alleles (i.e., allelic bias), as measured by MPRA (FDR < 5%). Bar plot shading corresponds to which promoter a variant was paired with when it demonstrated allelic bias (light shade: both, medium shade: *MYBPC2*, dark shade: SCP1). Total number of variants for each condition is listed above the corresponding bar. **(C)** UpSet plot shows counts of variants showing allelic bias in each subset of conditions (filled dots). **(D,E)** MPRA signals with log2(RNA/DNA) plotted vs alleles (Ref: reference, Alt: alternate) or a negative control, for **(D)** an example of allele-specific activity favoring the alternate allele primarily in differentiated conditions; and **(E)** a stimulation-specific example. (p-values in **(C)** and **(D)** from *post hoc* pairwise t-tests after passing mpralm 5% FDR threshold; N = 4 replicates per group)

We used mpralm (Myint et al. 2019) to estimate oligo activity in each individual condition (**Supplemental Tables 10-13**). We identified 303 active oligos in undifferentiated cells, 103 in basal differentiated cells, 38 in AICAR-stimulated cells, and 76 in palmitate-stimulated cells (FDR-adjusted *p* < 0.05; **Supplemental Figure 3A**). Across all four conditions, most active oligos were paired with the SCP1 promoter compared to *MYBPC2* or both promoters, ranging from 57.2% (n = 59/103) in the differentiated condition and 71.1% (n = 27/38) in the AICAR-stimulated condition (**Supplemental Table 14**). Within these oligo sets, we also examined the number of corresponding variants with at least one active allele and identified 208 unique variants in undifferentiated cells, 76 in differentiated cells, 28 in AICAR-stimulated cells, and 59 in palmitate-stimulated cells (**Supplemental Figure 4B**). Next, we compared active oligo sets across conditions which revealed a shared set of 21 active oligos, the majority of which were paired with the *MYBPC2* promoter (n = 20/21, 95.2%; **Supplemental Figure 4C**). The largest proportion of condition-specific active oligos was displayed by the undifferentiated group (276) with only 20, 3, and 3 uniquely active oligos in differentiated, palmitate-stimulated, and AICAR-stimulated groups, respectively.

We next tested for differential activity across alleles, focusing on variants where at least one allele was significantly active. (**Supplemental Figure 4B**, **Supplemental Tables 15-18**). We identified 128 variants with significant allelic bias in undifferentiated cells, 48 in differentiated cells, 18 in AICAR-stimulated cells, and 36 in palmitate-stimulated cells (FDR-adjusted *p* < 0.1; **Figure 3B**). As with the active oligo analysis, the majority of variants with allelic bias in each condition were paired with the SCP1 promoter, from 55.6% (n = 20/36) in palmitate-stimulated cells and 72.2% (n = 13/18) in AICAR-stimulated cells (**Supplemental Table 19**). We discovered a set of 7 variants with allelic bias across all four conditions, all of which displayed allelic bias when paired with the *MYBPC2* promoter. A separate set of 122 variants showed condition-specific allelic bias (**Figure 3C**). Variants that displayed allelic bias across more than one condition did so in the same direction in all cases except one (rs4782722 paired with the SCP1 promoter; **Supplemental Table 20**).

Given the broad effects of differentiation upon gene expression and MPRA activity, we sought to identify examples where allelic effects were also influenced by differentiation state. One such example is rs11867290, which displays significant allelic bias in both basal differentiated and palmitate-stimulated cells, with nearly significant allelic bias in AICAR-stimulated cells, where the alternate allele (G) displays higher enhancer activity than the reference allele (T) (**Figure 3D**; *p* = 0.034, 0.132, and 0.017 in differentiated, AICAR-stimulated, and palmitate-stimulated cells, respectively; *post hoc* pairwise t-tests). By contrast, both alleles are active in undifferentiated cells, but without significant allelic bias. rs11867290 is located in the second intron of *MARCHF10* and is a lead signal variant for height (Yengo et al. 2022), where the T allele is associated with decreased height. This variant overlaps an ENCODE-annotated enhancer for several metabolic tissues, including primary skeletal muscle and skeletal muscle-derived myoblasts and myotubes (ENCODE Project Consortium 2012; ENCODE Project Consortium et al. 2020; Luo et al. 2020).

We also identified variants that displayed allelic bias in a more restricted pattern, including rs34003091 (**Figure 3E**). While this variant enhances MPRA activity in both undifferentiated and differentiated cells, it displays allelic bias in neither condition. By contrast, the reference allele (T) is more active than the alternate allele (C) in both AICAR- and palmitate-stimulated cells (*p* = 0.01 and 0.017 in AICAR- and palmitate-stimulated, respectively; *post hoc* pairwise t-tests). rs34003091 is a lead signal variant for total cholesterol (Graham et al. 2021) and in high linkage with lead variants for LDL cholesterol (R^2^ = 1 with rs35081008 and R^2^ > 0.9 with rs34503352 across 1000 Genomes populations; Klimentidis et al. 2020; Sakaue et al. 2021; Hoffmann et al. 2018). The T allele is associated with decreased total cholesterol and on a shared haplotype with LDL cholesterol-raising alleles. This variant is located in a chromHMM-predicted weak promoter for *ZNF329* in HepG2 hepatocytes and also colocalizes with e- and caQTLs for the same gene in liver tissue (Currin et al. 2021; Etheridge et al. 2020). The authors of this study also used a luciferase assay in HepG2 hepatocytes to evaluate enhancer activity at this variant and observed allelic effects with concordant direction of effect to what we observed here (Pandey et al. 2024). Similar to the liver, skeletal muscle is an important regulator of lipid metabolism. One previous study observed decreased *ZNF329* expression in skeletal muscle biopsies taken from obese individuals before and after bariatric surgery, indicating that skeletal muscle may play a role in trait variation at this locus (Gancheva et al. 2019).

### Regulatory features and condition-specific activity at metabolic trait-associated variants

We next set out to identify upstream transcriptional regulators which may underlie the allelic effects observed in our MPRA. We focused on rs490972, a lead variant for serum triglyceride levels (DeForest et al. 2024) and in tight linkage with a separate lead variant (rs1625595; R^2^ > 0.97 across 1000 Genomes populations) for both metabolic syndrome (Park et al. 2024) and self-reported moderate-to-vigorous physical activity (Wang et al. 2022). In most of the conditions assayed by MPRA, this variant displayed strong allelic bias with the alternate allele (A) showing lower activity in comparison to the reference allele (G) (**Figure 4A**; *p* = 0.003, 0.004, and 0.093 in undifferentiated, differentiated, and palmitate-stimulated cells, respectively; *post hoc* pairwise t-tests). rs490972 overlaps several motifs for SP family transcription factors, including SP1 and SP4 and the reference allele (G) of this variant matches the consensus base at a high-information content position (**Figure 4B**). The specificity protein (SP) family of transcription factors are zinc-finger proteins that play key roles in transcriptional regulation of foundational cellular functions, such as growth-regulation and developmental processes (Kaczynski et al. 2003), and recognize and bind to GC-rich promoter sites to regulate cellular function. To nominate which of these two TFs may underlie the allelic difference in activity at this site, we compared their mRNA expression with the MPRA activity of the motif-preserving reference allele. We observed significant positive correlation (Spearman R^2^ = 0.761; *p* = 0.002) only for *SP1* but not *SP4* (**Figure 4C**). We predict that the alternate allele A disrupts the SP motif leading to decreased binding and reporter activity which is further exaggerated when *SP1* expression is decreased. This aligns with the observed reporter activity where this variant is only an enhancer in the undifferentiated group and is neither active nor displays allelic bias in the AICAR-stimulated group. Together, these data suggest that regulatory activity at the metabolic trait-associated variant rs490972 may be mediated by SP binding, restricted to conditions where the appropriate TF is expressed.

**Figure 4.**
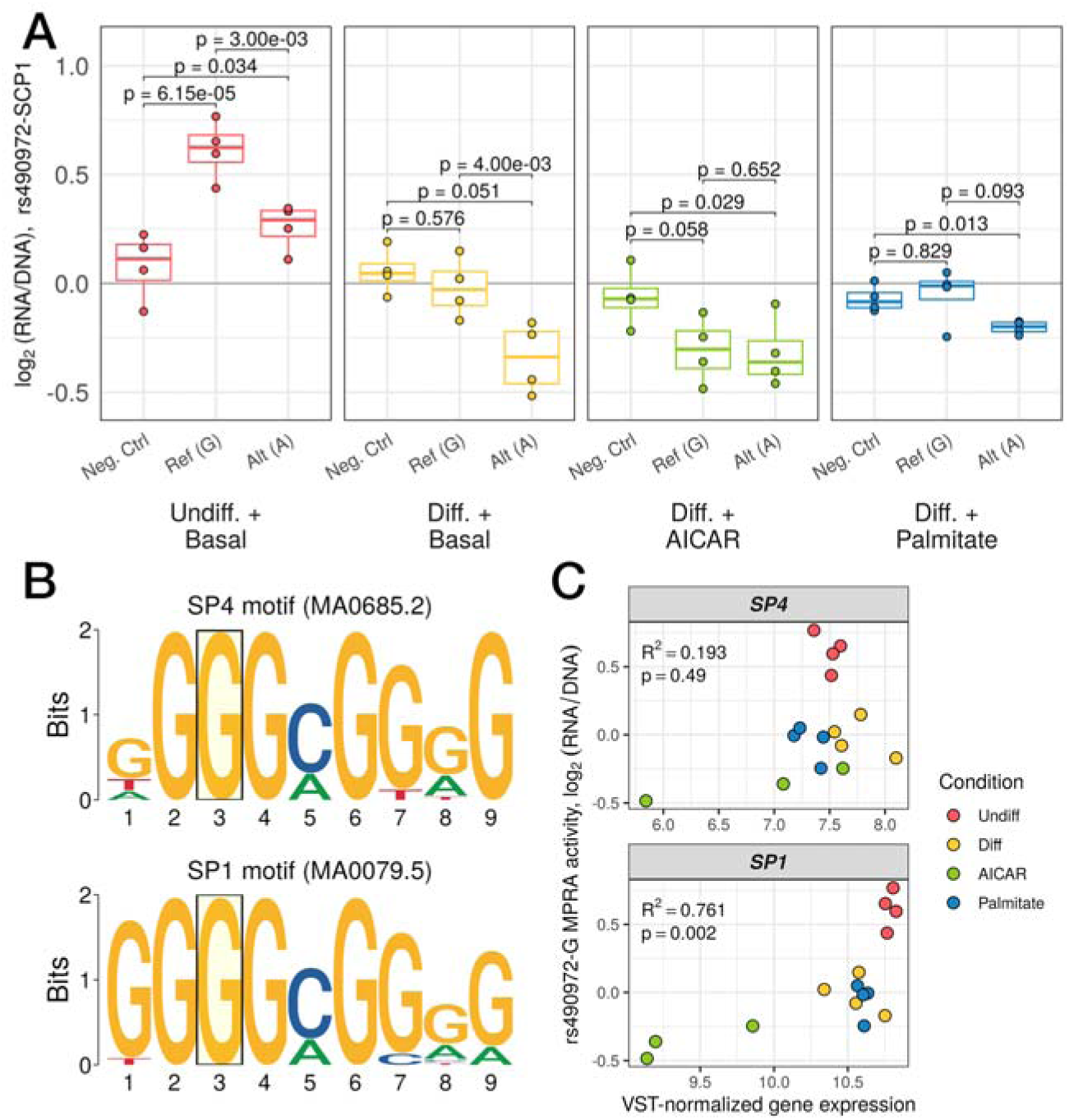
**(A)** Regulatory activity of a negative control sequence compared to the reference, G, and alternate, A, alleles of rs490972. Each point represents a single replicate and groups are color-coded (p-values from *post hoc* pairwise t-tests after passing mpralm 5% FDR threshold). **(B)** Logo plots of the motifs for SP4 (top) and SP1 (bottom). The position corresponding to where rs490972 overlaps both motifs is highlighted by a transparent yellow box. The reference allele of this variant, G, is the consensus nucleotide for both motifs. **(C)** Scatterplots of variance stabilizing transformation-normalized RNA expression counts for *SP4* (top) and *SP1* (bottom) compared to log_2_(RNA/DNA) for the reference allele of rs490972. Spearman correlation coefficient and p-value are displayed in the top left corner for each comparison. (N = 3 replicates for AICAR-stimulated group, 4 replicates for other three groups for RNA-seq data; N = 4 replicates per group for MPRA data)

## Discussion

In this study, we examined how transcriptional regulation and gene expression respond to differentiation and stimulation, and how those effects interact with allelic variation in a model of human skeletal muscle. We focused on skeletal muscle due to its involvement in metabolic disease processes, and investigated both developmental and environmental state differences using four different conditions. We differentiated LHCN-M2 proliferating myoblasts into terminally differentiated myotubes to assess developmental state differences, and we perturbed differentiated cells with either AICAR to mimic exercise, or palmitate to induce insulin resistance, models for different physiologically relevant cellular states.

A wide swath of genes showed differential expression between the conditions tested including the key muscle fate determination gene *MYF5,* which was induced during differentiation. AICAR treatment was marked by cellular responses to oxidative stress, while shifts after palmitate treatment included pathways involved in lipid metabolism. Enrichment for gene sets and pathways related to each of the chosen perturbations confirms that these treatments are appropriate models of the dynamic changes in state that muscle cells undergo during differentiation and disease processes.

Concurrently, we used an allelic MPRA to test metabolic-disease associated variants across the same four stimulatory states. Remarkably, clustering cells based on the MPRA measurements across 735 oligos recapitulated a qualitatively similar arrangement to PCA performed using whole-transcriptome RNA-seq, demonstrating that the selected conditions’ regulatory effects can be read out by MPRA. We noted that undifferentiated cells displayed the highest level of regulatory activity followed by differentiated cells in basal media, palmitate- and AICAR-stimulated cells. The basis for these differences is not clear, but they suggest that the undifferentiated, proliferative state may be globally more transcriptionally permissive, similar to what is observed during retinoic acid-induced differentiation of embryonic stem cells (Bulut-Karslioglu et al. 2018).

We also highlighted several variants in which our MPRA results demonstrated condition-specific allelic bias. One variant, rs11867290, displayed condition-specific effects reflecting differentiation state changes. This variant was recently uncovered as a lead signal variant for height (Yengo et al. 2022). rs11867290 demonstrated allelic bias in all conditions except for the basal undifferentiated cells. Another variant associated with total and LDL cholesterol (Graham et al. 2021; Klimentidis et al. 2020; Sakaue et al. 2021), rs34003091, only displayed allelic bias in AICAR- and palmitate-stimulated cells, illustrating a potential condition-specific effect. These instances of differentiation-state and environmental-induced differences on regulatory activity indicate that the function of regulatory variants are influenced by dynamic environmental changes and prompts further investigation of context-specific activity.

Lastly, to explore potential regulatory mechanisms functioning at disease-associated variants from our MPRA analysis, we investigated a specific variant, rs490972, which overlapped motifs for the SP family transcription factors SP1 and SP4. We hypothesize that regulatory activity at this loci may be mediated by SP1 transcription factor binding based on the differential regulatory activity between conditions. Taken together, these results reinforce the importance of examining cellular environments when assessing the functional impact of regulatory variation. While other studies have explored the effect of stimulatory states on regulatory variation as assessed in an MPRA, such as in pancreatic β cells through investigating steady state and endoplasmic reticulum stress conditions (Khetan et al. 2021), or at different time-points of neural differentiation (Kreimer et al. 2022), this study is the first that we are aware of to employ this framework in a model of skeletal muscle. As the primary site for insulinLstimulated glucose uptake, skeletal muscle is an essential tissue in which to understand how regulatory variants affect metabolic traits. We explored regulatory activity across a small panel of variants, many of which were selected based on prior evidence of regulatory activity in a luciferase assay. Our results serve as a proof of principle for scaling up MPRAs to examine the thousands of variants across hundreds of loci linked to metabolic diseases, as well as saturation-scale efforts to mutationally dissect individual loci. Moreover, while the use of an MPRA library provided robust initial findings, investigating these variants in their native genomic contexts (e.g., via CRISPR editing) is a crucial next step to fully characterize and understand their function. Finally, the regulatory inferences presented here lay the foundation for future work that integrates other functional genomic datasets to refine the regulatory mechanisms driving activity at disease-associated loci. For instance, ChIP-seq for specific transcription factors of interest (e.g., SP1, SP4) in LHCN-M2 cells across different perturbation states could provide further evidence for involvement at variants of interest.

Altogether, this study builds upon the current understanding of how metabolic disease-associated variants are regulated in skeletal muscle across developmental state changes and in response to relevant perturbations. Our findings provide evidence for widespread context-specific regulatory effects at metabolic disease-associated loci. Future studies should take this into consideration and assay an even broader set of tissues and contexts to eventually develop a comprehensive understanding of disease mechanisms.

## Methods

### Cell culture

We obtained LHCN-M2 human skeletal muscle myoblasts from Evercyte. We maintained LHCN-M2 cells on 0.1% gelatin-coated flasks in 4:1 DMEM (4.5 g/L glucose, glutamine, bicarbonate):Medium 199 (with bicarbonate) containing 15% FBS, 20 mM HEPES, 30 ng/mL zinc sulfate, 1.4 ug/mL vitamin B12, 55 ng/mL dexamethasone, 2.5 ng/mL recombinant human hepatocyte growth factor, 10 ng/mL basic fibroblast growth factor, and 600 U/mL penicillin/streptomycin. To differentiate the LHCN-M2 myoblasts to myotubes, we maintained cells in LHCN-M2 differentiation media containing 50:1 DMEM (1 g/L glucose):heat-inactivated horse serum for 7 days with daily media changes.

### Insulin stimulated glucose uptake assay

Approximately two hours prior to applying perturbations, we exchanged LHCN-M2 media for unsupplemented DMEM (1 g/L glucose). Then, we incubated cells in basal culture media, basal differentiation media, or differentiation media supplemented with 2mM AICAR or 0.5mM palmitate for 24 hours followed by glucose starvation for 3.5 hours in modified Krebs Ringer buffer with BSA (pH 7.2) containing 1X SAB stock (10X SAB: 1.14M NaCl, 47mM KCl, 12mM KH_2_PO_4_, 11.6 mM MgSO_4_), 1M HEPES, 25 mM CaCl_2_, 0.2% BSA (w/v), and 25.5 mM NaHCO_3_. Afterward, we added fresh Krebs Ringer buffer with or without 100 nM insulin for 30 minutes. We measured 2-deoxyglucose uptake with the Glucose Uptake-Glo™ Assay and assessed cell viability in parallel using the CellTiter-Glo® Luminescent Cell Viability assay, both according to the manufacturer’s instructions (Promega, Fitchburg, WI, USA). We used a GloMax Multi+ Detection System (Promega, Fitchburg, WI, USA) to measure luminescence intensity (relative light units, RLU). We normalized raw data by subtracting an average of background RLU from all empty wells. We further normalized glucose uptake values using CellTiter-Glo measurements to account for variation in cell number and viability.

### MPRA library design, construction, and delivery

We designed, constructed and delivered the MPRA library as described in Tovar et al. 2023 based on methods in Tewhey et al. 2016 and Gordon et al. 2020. We gathered signal variants relevant to T2D and metabolic traits and designed 198-bp oligo sequences. We added 16-bp flanking adapter sequences for PCR amplification and cloning, and obtained a pool of 1,255 full-length 230-bp oligos from IDT. We added barcodes and a promoter cloning scaffold via PCR, and cloned amplified oligos into a modified pMPRA1 vector (a gift from Tarjei Mikkelsen; Addgene #49349) using Golden Gate assembly with PaqCI. We digested this assembly with SfiI to remove empty backbones then transformed into electrocompetent 10-beta cells. We inserted clonal promoter fragments using Golden Gate assembly and digested with AsiSI to remove promoterless constructs. We transformed this final assembly into electrocompetent 10-beta cells, expanded in 150 mL cultures, and isolated plasmid using the ZymoPURE Plasmid Maxiprep Kit.

### MPRA barcode pairing

To create barcode-oligo pairs, we sequenced the promoterless constructs. We generated initial amplicons with primers specific to the regions immediately 5’ of the oligo and 3’ of the barcode. We amplified these pairing amplicon using P7 and P5 stubs, then added dual sequencing indexes. We sequenced the resulting libraries on an Illumina NovaSeq 6000 and received paired-end 150-bp reads.

### MPRA/RNA-seq sample generation

To deliver the MPRA library to the LHCN-M2 human skeletal myoblasts, we ported the assembled MPRA block (oligo, barcode, promoter, GFP) to a lentiviral transfer vector (a gift from Nadav Ahituv; Addgene #137725) via restriction cloning (Gordon et al. 2020; Inoue et al. 2017). This lentiviral transfer vector was used by the University of Michigan viral vector core to produce infectious lentiviral particles. The undifferentiated cells were maintained and passaged while other samples per condition were differentiated for 5 days. On day 6, we infected 15 x 10^6^ LHCN-M2 human skeletal myoblasts per replicate with our MPRA library at an MOI of ∼10. On day 7, we exchanged media for fresh basal media (undifferentiated) or differentiation media and incubated cells for 24 hours. Subsequently, cells were incubated in basal media, basal differentiation media, or differentiation media supplemented with 2mM AICAR or 0.5mM palmitate for 24 hours. We isolated RNA and gDNA from each replicate using the Qiagen AllPrep DNA/RNA mini kit.

### RNA-seq library generation

We used the NEBNext® Poly(A) mRNA Magnetic Isolation Module (New England BioLabs, Ipswich, MA) to enrich poly(A)+ RNA transcripts from total RNA for RNA library preparation and sequencing. Afterward, we used the NEBNext® Ultra™ II Directional RNA Library Prep kit (New England BioLabs, Ipswich, MA) to prepare sequencing libraries following manufacturer’s instructions. At this stage, we excluded one sample (ATS0272, AICAR replicate 2) due to low library concentration. We sequenced libraries on an Illumina NovaSeqX and received paired-end 150-bp reads.

### MPRA barcode library generation

After gDNA and RNA collection, we treated RNA with the TURBO DNA-free™ kit (Thermo Fisher Scientific, MA, USA) following the manufacturer’s instructions. We synthesized cDNA from the resulting cleaned RNA using the SuperScript™ III First-Strand Synthesis System (Thermo Fisher Scientific, MA, USA) with a GFP-specific primer containing a 15-bp UMI, and subsequently amplified cDNA via PCR. We attached sequencing adapters and a 13-bp UMI by PCR, then amplified libraries using P7 and P5 stubs. We added dual sequencing indexes to both cDNA and gDNA libraries in a final 6-cycle PCR reaction. We sequenced the resulting cDNA/gDNA libraries on an Illumina NovaSeq 6000 and received paired-end 150-bp reads.

### RNA-seq data analysis

RNA-seq raw data was processed with a custom pipeline. We mapped reads to hg38 using STAR (v2.7.9a) (Dobin et al. 2013). We used SAMtools to filter high-quality read pairs (-q 255 -F 4 -F 256 -F 2048; v. 1.9 using htslib v. 1.9) (Li et al. 2009). We used FastQC to assess the quality of raw sequence data (v. 0.11.5) and MultiQC to summarize quality results (v. 1.11) (Andrews 2010; Ewels et al. 2016). To perform quality control on aligned reads, we used QoRTs (Quality of RNA-seq Tool-Set) (v. 1.3.6) (Hartley and Mullikin 2015). Given the dominant effects of differentiation state on gene expression (see **Figure 2A**), we chose to split the data into two subsets for differential expression analysis: (1) basal differentiated versus basal undifferentiated samples and (2) basal differentiated versus differentiated with AICAR or differentiated with palmitate samples. We conducted differential expression analysis using DESeq2 v.1.34.0 (Love et al. 2014) using a log_2_ fold change cutoff of 1 and a 5% false discovery rate. To perform pathway enrichment analysis, we split sets of differentially expressed genes based on log_2_FC to assign to each condition. For example, for the basal undifferentiated versus basal differentiated contrast, we assigned genes with a positive log_2_FC to the basal differentiated state and vice versa for the basal undifferentiated state. We then used g:Profiler v.0.2.3 (Raudvere et al. 2019) to perform pathway enrichment, subset results to the “Biological Process” and “Molecular Function” ontologies, and used rrvgo (Sayols 2023) with a threshold of 0.7 to cluster similar terms. To improve interpretability, we removed terms with fewer than 10 and more than 1000 member genes.

### MPRA data analysis

To create the barcode-oligo pairing dictionary and process MPRA sequencing data, we used custom pipelines as described previously (Tovar et al. 2023). We used the R package mpra to estimate oligo activity and allelic bias (Myint et al. 2019). For our activity analysis, we used a 5% FDR threshold. For our allelic bias analysis, we required that at least one allele for a variant was active and to identify variants with differential allelic activity, we used a 10% FDR threshold.

### Data and code availability

Primer sequences used for MPRA library cloning and sequencing library generation are available in **Supplemental Table 21**. All processed RNA-seq counts and MPRA data are accessible through the Gene Expression Omnibus (GSE282191). Raw sequencing data are deposited on the NCBI Sequence Read Archive and accession numbers are listed in the GEO records. Complete results files from both sets of analyses are available as Supplemental Tables referenced in this article.

All code used to preprocess and analyze data presented here are available on Github. We used custom preprocessing pipelines for RNA-seq data (https://github.com/porchard/RNAseq-NextFlow) and MPRA barcode counting data (https://github.com/adelaidetovar/MPRA-Nextflow). Scripts used to analyze and visualize processed data are available as a separate repository at https://github.com/adelaidetovar/lhcn-perturb-mpra.

## Acknowledgements

The authors would like to acknowledge members of the Kitzman and Parker labs (University of Michigan) for their critical feedback. The authors would also like to thank the University of Michigan Viral Vector Core for producing lentivirus.

## Contributions

Conceptualization by A.T., J.O.K., and S.C.J.P. Oligo design and MPRA library cloning by K.N. Data acquisition by K.N. and G.B. Data processing and analysis by K.N., G.B., and A.T. K.N. wrote the manuscript with input from and editing by A.T., G.B., J.O.K., and S.C.J.P.

