## Supplemental Figures for "Functional dissection of metabolic trait-associated gene regulation in steady state and stimulated human skeletal muscle cells"

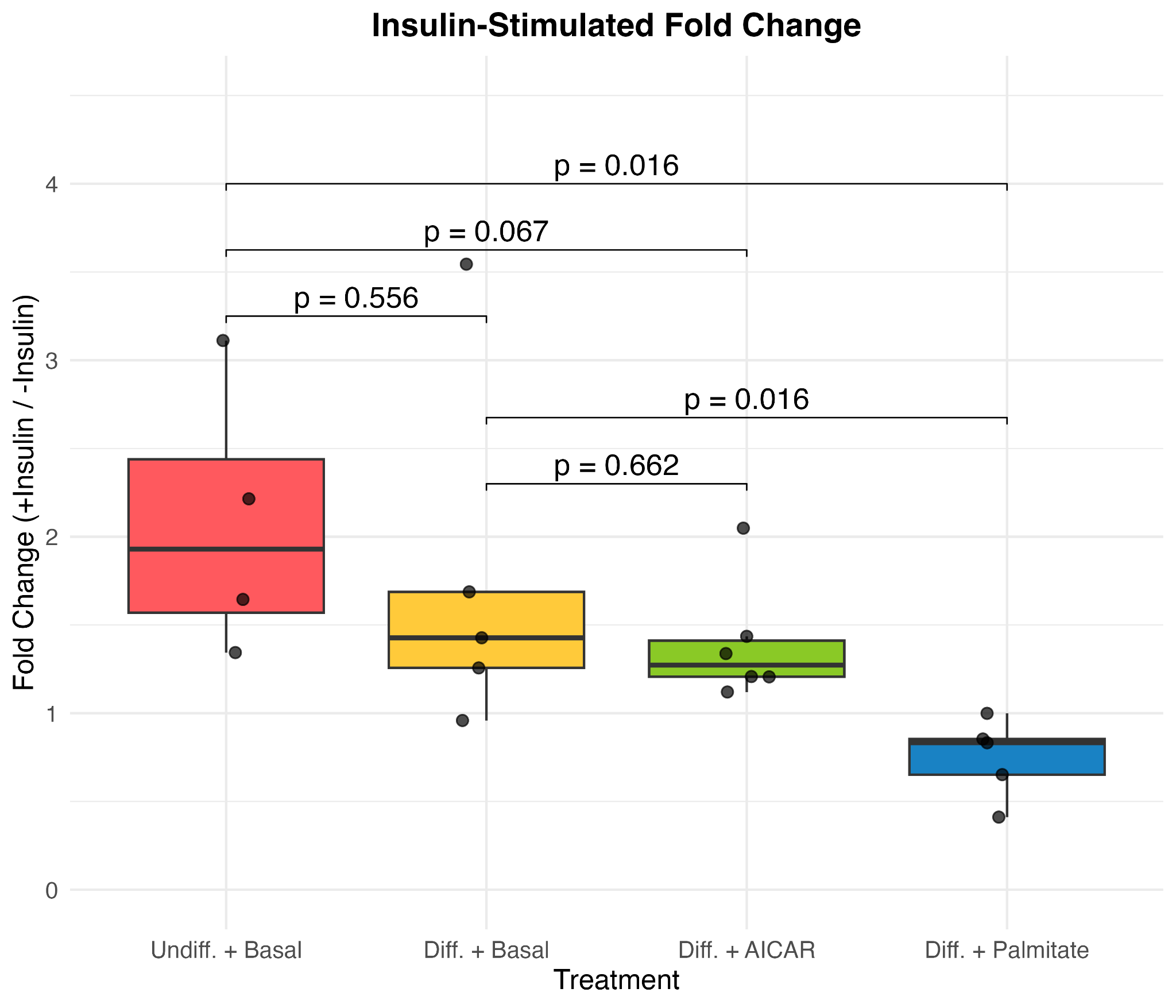

**Supplemental Figure 1. Insulin-stimulated glucose uptake across conditions, reflecting metabolic changes.** Box plots displaying fold-change of insulin-stimulated glucose uptake relative to paired replicates in the absence of insulin stimulation, measured as normalized luminescence intensity. Each point corresponds to the ratio for an individual replicate, and box colors indicate condition. (p-values from Wilcoxon signed-rank test; n = 4 for undifferentiated cells; n = 5 for differentiated + basal media and differentiated + palmitate stimulation; n = 6 for differentiated + AICAR treatment)

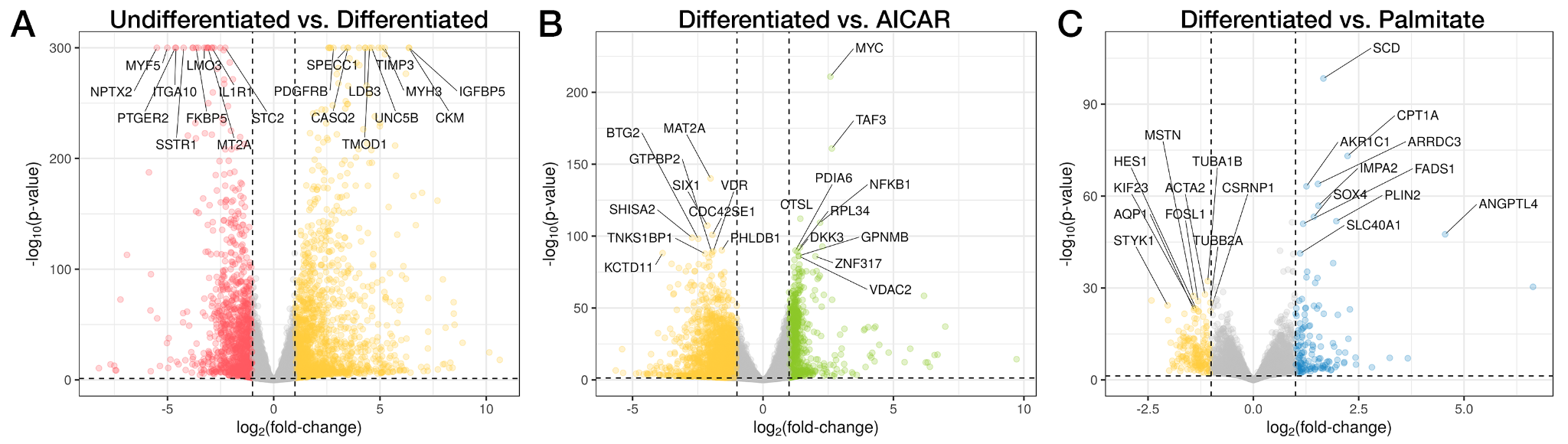

**Supplemental Figure 2. Distinct sets of genes are differentially expressed across stimulation conditions.** Volcano plots displaying differential expression compared across **(A)** undifferentiated (red, left) and differentiated (yellow, right) groups; **(B)** differentiated (yellow, left) and AICAR-stimulated groups (green, right); and **(C)** differentiated (yellow, left) and palmitate-stimulated groups (blue, right). Vertical dashed lines correspond to log_2_FC = -1 and 1. Fold-change estimates obtained via DESeq2 (multiple testing corrected at FDR < 5%). Horizontal dashed line corresponds to FDR-adjusted *p* = 0.05. The top ten downregulated and upregulated genes are labeled in each volcano plot. (N = 3 replicates for AICAR-stimulated group, 4 replicates for other three groups for RNA-seq data; n = 15,285 genes for undifferentiated vs. differentiated comparison and n = 15,677 genes for differentiated vs. AICAR or palmitate comparisons)

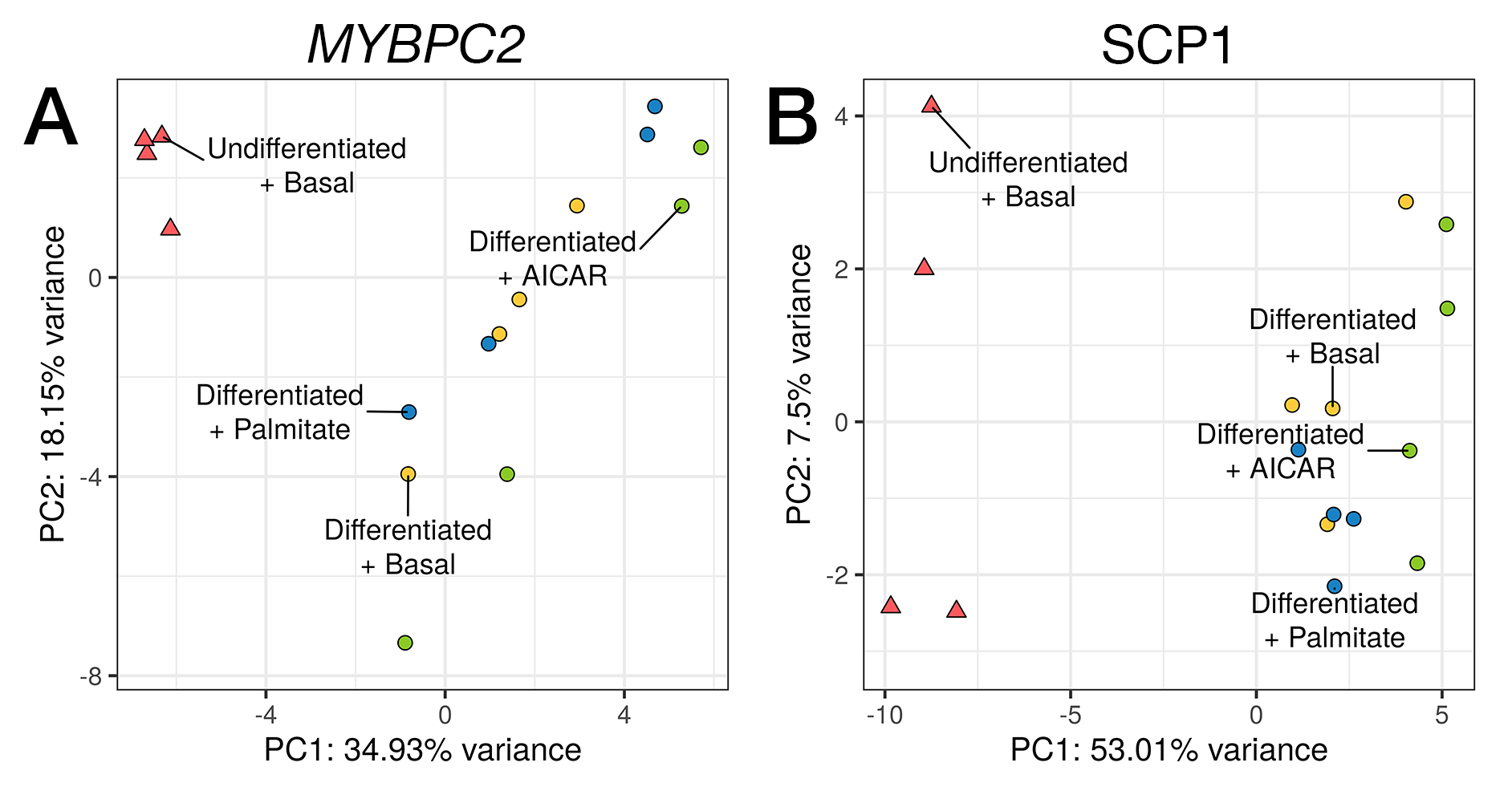

**Supplemental Figure 3. Principal components analysis of MPRA activity for (A) *MYBPC2* and (B) SCP1 promoter.** Principal components analysis (PCA) of regulatory activity expressed as log_2_(RNA/DNA) for all oligos. Point color corresponds to group and point shape corresponds to differentiation status. (n = 4 per condition)

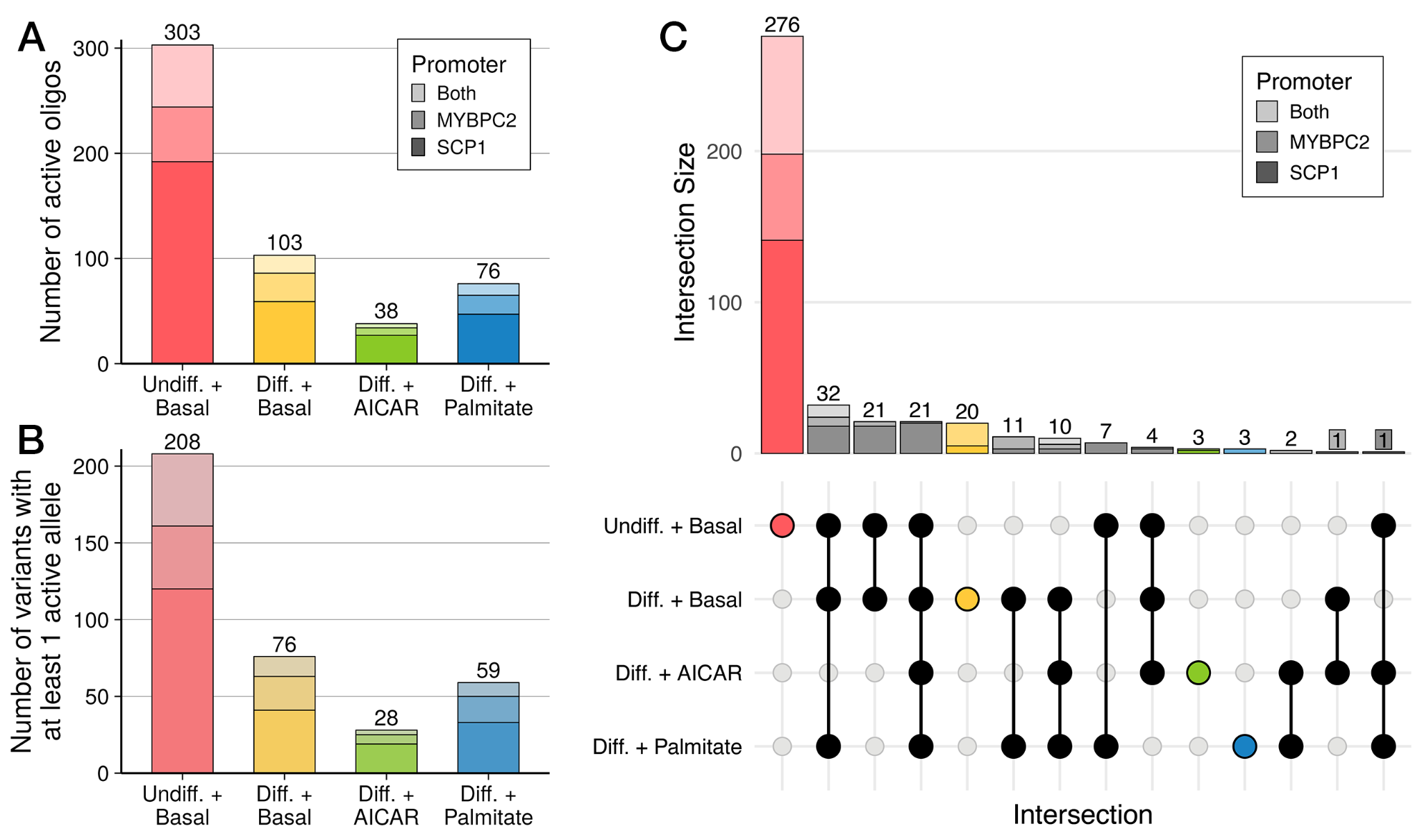

**Supplemental Figure 4. Undifferentiated myoblasts display distinct regulatory patterns compared to other three groups.** **(A)** Stacked bar plot displaying the number of active oligos, as measured by MPRA (FDR < 5%). Bar plot shading corresponds to which promoter an oligo was paired with when it was active (light shade: both, medium shade: *MYBPC2*, dark shade: SCP1). Total number of oligos for each condition is listed above the corresponding bar. **(B)** Stacked bar plot displaying the number of variants with one or more active alleles in each condition. **(C)** UpSet plot exhibiting sets of oligos based on whether they were active in one or more conditions. The number of variants in each shared or unique set is displayed in the top stacked bar plot with the same shading as described in **(A)**. The matrix plot on the bottom indicates which condition(s) the bars correspond to. (n = 4 per condition)
